# Latent evolution of biofilm formation depends on life-history and genetic background

**DOI:** 10.1101/2023.04.06.535679

**Authors:** Amandine Nucci, Eduardo P.C. Rocha, Olaya Rendueles

**Affiliations:** Institut Pasteur, Université de Paris Cité, CNRS, UMR3525, Microbial Evolutionary Genomics, F-75015, Paris, France

**Keywords:** latent phenotypic evolution, chaperon-usher system Running title: Mutations in *mrkD* alter biofilm formation

## Abstract

Adaptation to one environment can often generate phenotypic and genotypic changes which impact the future ability of an organism to thrive in other environmental conditions. In the context of host-microbe interactions, biofilm formation can increase survival rates *in vivo* upon exposure to stresses, like the host’s immune system or antibiotic therapy. However, how the generic process of adaptation impacts the ability to form biofilm and how it may change through time has seldomly been studied. To do so, we used a previous evolution experiment with three strains of the *Klebsiella pneumoniae* species complex, in which we did not specifically select for biofilm formation. We observed that changes in the ability to form biofilm happened very fast at first and afterwards reverted to ancestral levels in many populations. Biofilm changes were associated to phenotypic changes in population yield and surface polysaccharide production. Genotypically, mutational targets in the tip adhesin of type III fimbriae (*mrkD*) or the *fim* switch of type I fimbriae were driven by nutrient availability during evolution, and their impact on biofilm formation was dependent on capsule production. Analyses of natural isolates revealed similar mutations in *mrkD*, suggesting that they also play an important role in adaptation outside the laboratory. Our work reveals that the latent evolution of biofilm formation, and its evolutionary dynamics, depend on nutrient availability, the genetic background and other intertwined phenotypic and genotypic changes. Ultimately, it suggests that small differences in the environment can alter an organism’s fate in more complex niches like the host.

## INTRODUCTION

One of the central questions in microbial evolutionary biology is understanding the mechanisms by which bacteria expand their ecological breadth. The niche shift hypothesis postulates that the process of adaptation to a different environment can result from rapid adaptive changes via mutation or horizontal gene transfer [1, 2], leading to diversification and opening the possibility of exploiting novel niches [3]. Bacteria may have to contend with novel stresses to adapt. This may often involve forming a biofilm, which generically increases tolerance to a broad range of stresses [4]. Such resilient surface-attached multicellular structures are ubiquitous and the prevalent prokaryotic lifestyle [5].

In the context of host-microbe interactions, it has been shown that increased capacity to form biofilm correlates with the capacity to replicate and colonize multiple hosts, whereas bacteria with narrow host ranges are usually poor biofilm-formers [6, 7]. Within a host, biofilm formation offers numerous specific advantages, such as higher resistance to antimicrobials [8] and to antibody-mediated killing and phagocytosis [9]. During competition with other members of the microbiome, it can also lead to niche exclusion of direct competitors [10].

*Klebsiella pneumoniae* is a metabolically versatile enterobacterium with a very large carbon and nitrogen core metabolism [11]. This may partly explain its ubiquity and ecological breadth [12–14]. It can adopt a free-living lifestyle in the soil or in the water but it is mostly studied in its host-associated form, colonizing plants, insects and mammals, including humans, where it can be a found as a gut commensal. Hypervirulent strains of *K. pneumoniae* cause community-acquired infections which may result in pyogenic liver abscesses, but most of *K. pneumoniae* infections are opportunistic and health-care associated. They typically require a precolonization of the gastrointestinal epithelia prior to infecting other body sites [15].

Several factors impact the ability of *K. pneumoniae* to form biofilm and colonize host tissue, most notably two chaperon-usher systems [16]: the type I fimbriae encoded by the *fimA-K* operon and the type III fimbriae encoded by the *mrkA-I* operon. The former has been shown to preferentially bind to mannose residues in *E. coli* but not in *K. pneumoniae* [17], whereas the latter has high affinity to collagen [18, 19] and mediates adhesion to abiotic surfaces [20]. *In silico* studies have predicted the existence of many other chaperon–usher systems that could have specific tropism or be expressed in response to specific environmental cues [21].

In addition to surface adhesins, another important factor determining biofilm formation in *K. pneumoniae* is the extracellular capsule [22–25] produced by most isolates [26]. On the one side, some studies revealed that the capsule can strongly inhibit biofilm formation by masking surface adhesins [24] or by altering surface physico-chemical properties and thus limiting surface attachment and inter-cellular interactions [27, 28]. On the other side, presence of some *Klebsiella* capsules has been shown to increase the formation of biofilm and be required for its maturation [22]. Thus, the role of the capsule in biofilm formation is convoluted and depends both on the physical interactions between the capsule and the environment [25], and the genetic interactions between the capsule locus and the rest of the genome [23].

Numerous studies have focused on how different microbes increase biofilm formation by positively selecting for this trait [29–31]. Yet, how it changes when it is not under strong selection, or just as a mere by-product of the generic processes of adaptation is not currently understood. Indeed, adaptation of a given population to different novel environments may impact its ability to adhere and form a biofilm. This can have important consequences, for instance, in host colonization or increased tolerance to antibiotics. To study how biofilm formation in evolving populations changes through time, we took advantage of a previous evolution study in which we evolved in parallel six populations from three different strains of the *K. pneumoniae* species complex, and their respective non-capsulated mutants (Figure S1) [32]. We did so in five different liquid static environments, including artificial sputum (ASM) and urine (AUM), varying in carrying capacity (nutrient availability). Here, we measured the evolution of biofilm formation to determine whether it latently changes when it is not specifically selected for. If it does, we wished to enquire if this evolutionary process takes place in a progressive manner or evolves by leaps. We hypothesize that changes in biofilm could be the result of changes in other phenotypic traits that were under strong selection in our evolution experiment and which are known to affect biofilm formation. We thus specifically tested for correlation in changes in population yield or surface-attached polysaccharide production (capsule or others), and changes in biofilm formation. To link phenotype with genotype, we investigated whether the changes in biofilm formation were contingent with the presence of mutations in the two main types of fimbrial adhesins. Taken together our work highlights how the generic process of adaptation to non-biotic structured environments may promote the ability of a bacteria to form biofilm, and thus potentially expand its niche from the free-living environment to host colonization.

## RESULTS

### Changes in biofilm formation occur fast and depend on nutrient availability in the environment

To understand how adaptation shapes the ability of a bacterium to form biofilm, we quantified the biofilm in all evolving populations at regular intervals during the evolution experiment (day 15, 45, 75 and 102 – *i.e.* 100, 300, 500 and 675 generations). Taking all populations together, there was a fast and significant increase in the biofilm formation capacity by day 15 of *ca* +36%, which continued until day 45 (*ca* +50%) (One-sample Wilcoxon Rank Sum test, difference from 1, P < 0.001). Afterwards, a significant decrease is observed, to end up with total increase of *ca* ∼27% at the end of the experiment (One-sample Wilcoxon Rank Sum test, difference from 1, P = 0.01). We observed a high degree of parallel convergent evolution across replicates of the same ancestral genotype in each environment (Figure S2A). Notwithstanding, there were large across-treatment differences between environments and genotypes (Figure S2A). For instance, in Kva 342 populations, little change is observed in biofilm formation in populations evolving in M02, a steady increase is observed in LB, and divergent evolutionary paths were observed across capsule genotypes in ASM.

We had previously observed that the presence or absence of the capsule drives the direction and magnitude of evolutionary change in endpoint populations [32], yet how it affects the evolutionary dynamics remained to be tested. Initial changes in biofilm formation were similar across both capsule genotypes (Figure 1A). However, we observed divergent evolution between non-capsulated and capsulated populations towards the end of the evolution experiment, as biofilm formation in the latter decreased almost to ancestral levels (Figure 1A). This was mostly observed in Kpn strains in environments with low carrying capacity, namely M02 and AUM (Figure S2A). We thus tested whether the nutrient availability of the environment influenced evolutionary dynamics. In nutrient-rich environments with high carrying capacities (ASM and LB), populations increased biofilm formation steadily through the duration of the experiment (Figure 1B). In environments with lower carrying capacities, there was a similar increase in the ability to form biofilm compared to those in nutrient-rich, but this reverted fast to ancestral values (Figure 1C). Multifactorial ANOVA revealed that changes in biofilm were strongly dependent on the interaction between capsule genotype and nutrient availability (F = 123.4, P < 0.001). Independently, nutrient availability also strongly affected the evolution of biofilm formation (F = 52.17, P < 0.001), but not the ancestral capsule genotype (F = 3.05, P = 0.08).

**Figure 1.**
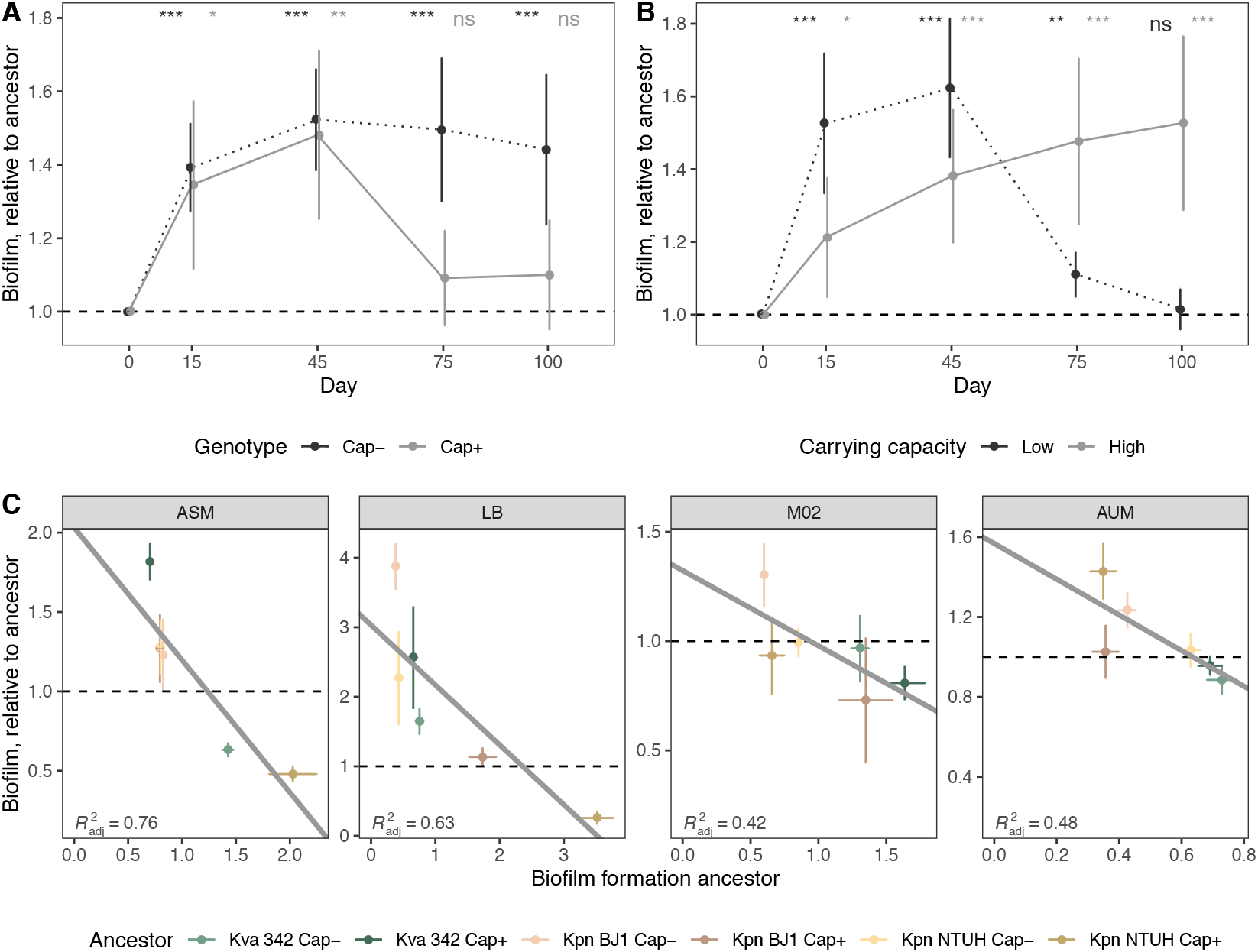
Changes in biofilm formation during ∼675 generations. **A.** Dynamics of biofilm formation in populations descending from capsulated (grey) or non-capsulated ancestor (dotted, black), relative to their respective ancestor, across all environments and strains. Grey points are shifted not to overlap with black points for visualization purposes. Error bars indicate interval of confidence (*α* = 0.05). Statistics represent One-sample Wilcoxon Rank Sum test, difference from 1. **P<0.01 and *** P<0.001. **B.** Dynamics of biofilm formation in populations evolving in high carrying capacity environments (ASM and LB, dotted black) compared to low carrying capacities (AUM and M02, in grey) relative to their respective ancestor, across all ancestral strains. **C.** Linear regression between the ancestral formation of biofilm per genotype (x-axis) and formation of biofilm after ∼675 generations (grey line). Each point represents the average of the independently evolving populations. Vertical error bars reflect the diversity of evolutionary outcomes (standard deviation from the mean). Horizontal error bars represent experimental variance of the phenotype (interval of confidence, 95%). R^2^ is indicated (P-values P<0.05, except for M02 where P=0.09).

We then tested how end-point populations had evolved as a function of the ancestral capacity of biofilm formation in each environment. We observed a strong negative correlation, across all environments, between the ancestral ability of each genotype to form biofilm and the relative change in biofilm formation (Figure 1C, P<0.05 for all except M02). The less biofilm the ancestor could produce, the larger the increase in biofilm formation observed at the end of the experiment. Inversely, genotypes that were already proficient biofilm formers tended to decreased biofilm production, for example capsulated Kpn NTUH in LB and ASM. This suggests that biofilm formation could be under stabilizing selection in *Klebsiella*.

Taken together, large changes in the capacity of forming biofilm happened during the first steps of adaptation. At longer evolutionary times, changes were mostly driven by nutrient availability.

### The environment determines how changes in population yield and surface polysaccharides influence biofilm formation

Adaptation to novel environments latently altered the ability of populations to form biofilm. Yet, our methodology to evaluate biofilm, *i.e.* by staining the extracellular matrix and attached cells with crystal violet, could be influenced by other variables, such as the changes in total population yield or in the production of surface polysaccharides. We tested how these two traits evolved throughout our evolution experiment (Figure S2BC, Table S1). Unfortunately, population yield in some capsulated populations could not be tested due to the emergence of hypermucoviscosity, which precludes accurate CFU assessment [32]. On average, evolving populations significantly increased yield (x̄ = +60%, One-sample Wilcoxon Rank Sum test, P< 0.001) (Figure S2B, Table S1), but as observed for biofilm formation, most changes occurred early during the evolution experiment. Similarly, surface polysaccharide production also increased (x̄ = +30%, One-sample Wilcoxon Rank Sum test, P= 0.0001), but only in environments with high carrying capacity (Figure S2C, Table S1).

To specifically test how changes in either yield or surface polysaccharides could influence biofilm formation, we correlated the degree of change relative to the ancestor of each of these two variables and the degree of change in biofilm formation. Despite the differences across ancestral genotypes, both yield and surface polysaccharides influenced changes in biofilm formation. However, these changes were different across environments and strongly depend on the carrying capacity of each environment (Figure 2). In environments with high carrying capacity, we observed a positive correlation between both yield and surface polysaccharides with biofilm formation. But in environments with low carrying capacity (AUM and M02), biofilm formation negatively correlated with changes in yield and surface polysaccharides. Despite such general trends, there are many exceptions. For instance, Kva 342 populations evolving in ASM increased population yield, yet biofilm formation was reduced. The correlations explain only a small fraction of the observed changes in biofilm formation, as indicated by their low *rho* values.

**Figure 2.**
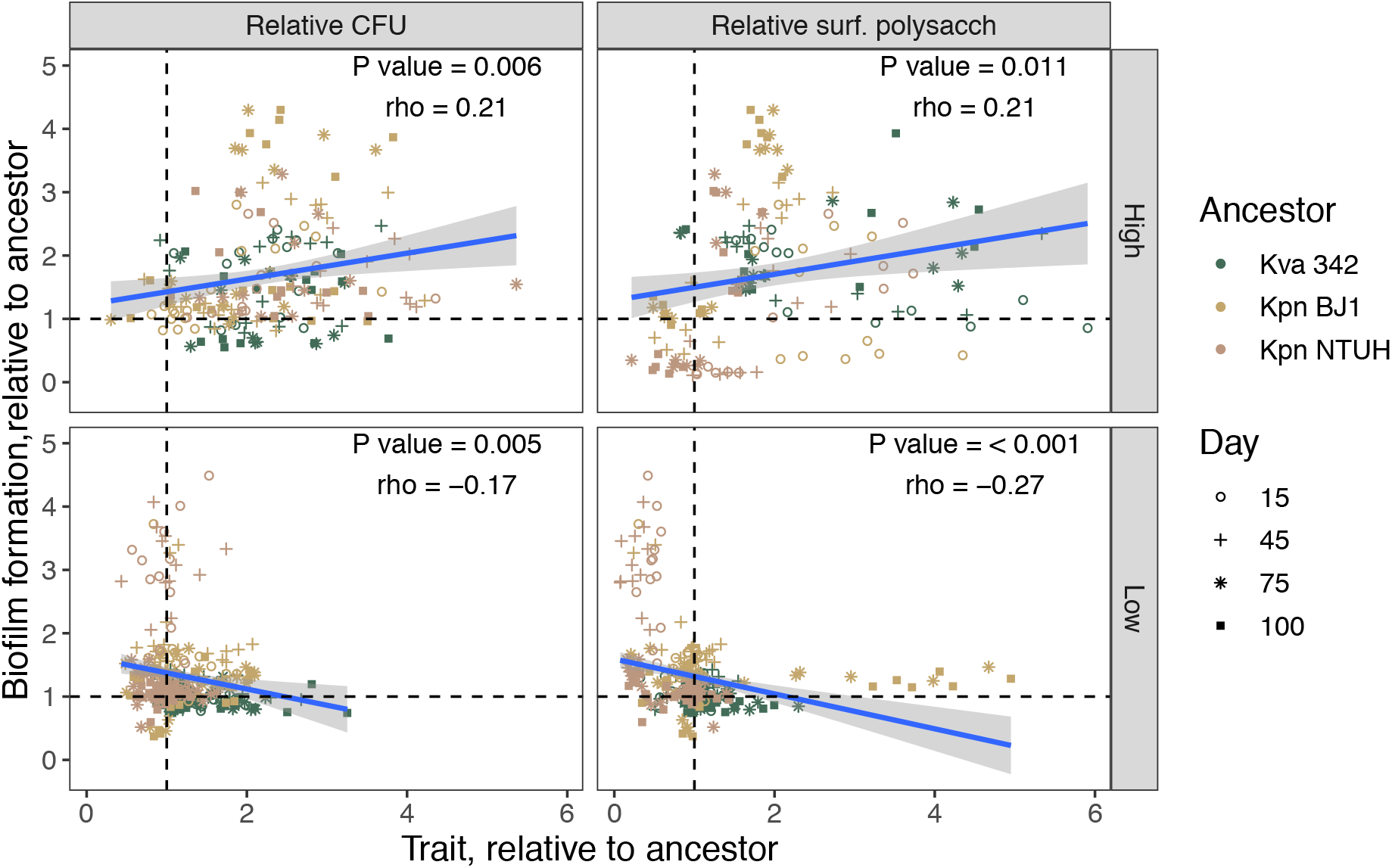
Correlation between changes in biofilm formation and co-evolving traits. Each shape corresponds to different time during the evolution experiment, and colors indicate different ancestral genotypes. Each dot is the average of at least three independent biological replicates. P-values correspond to a Spearman correlation test.

Overall, changes in adaptive traits like population yield and surface polysaccharide production correlated with changes in a latent phenotype, *i.e*. biofilm formation., but did not fully explain changes in the latter and depended on the environment to which populations were adapting.

### Convergent adaptation in *mrk* operon

To understand the genotypic factors involved in changes in biofilm formation, we analysed the genomic sequences of one randomly chosen clone from each population. We hypothesized that the convergent adaptation by repeated mutations in the *mrk* locus, which encodes for a major adhesin, could be largely responsible for changes in biofilm formation (Table S2). Indeed, our results show that populations in which mutations were detected in *mrkD*, the tip adhesin, but not in the rest of the operon, were associated with increased biofilm formation (Figure 3A). Mutations in *mrkD* tend to accumulate in populations evolving in rich environments (56% population in ASM or LB vs only 9% of populations in AUM or M02) (Figure 3B and Figure S3). This fits our observation that populations evolving in environments with high carrying capacities form more biofilm. Despite a lower mutational supply in populations evolving in low-nutrient environments, changes in biofilm in these populations can be observed after just fifteen days. These may indicate that such changes are due to other mutations (Figure 1A and 2). Of note, most mutations accumulate in the lectin binding domain (89%) which determines the binding specificity of the pili, in this case to type V collagen, as opposed to mutations in the pilin domain (11%), which would mostly influence the structure. Thus, this suggests that most mutations could be affecting surface affinity (Figure 3D).

**Figure 3.**
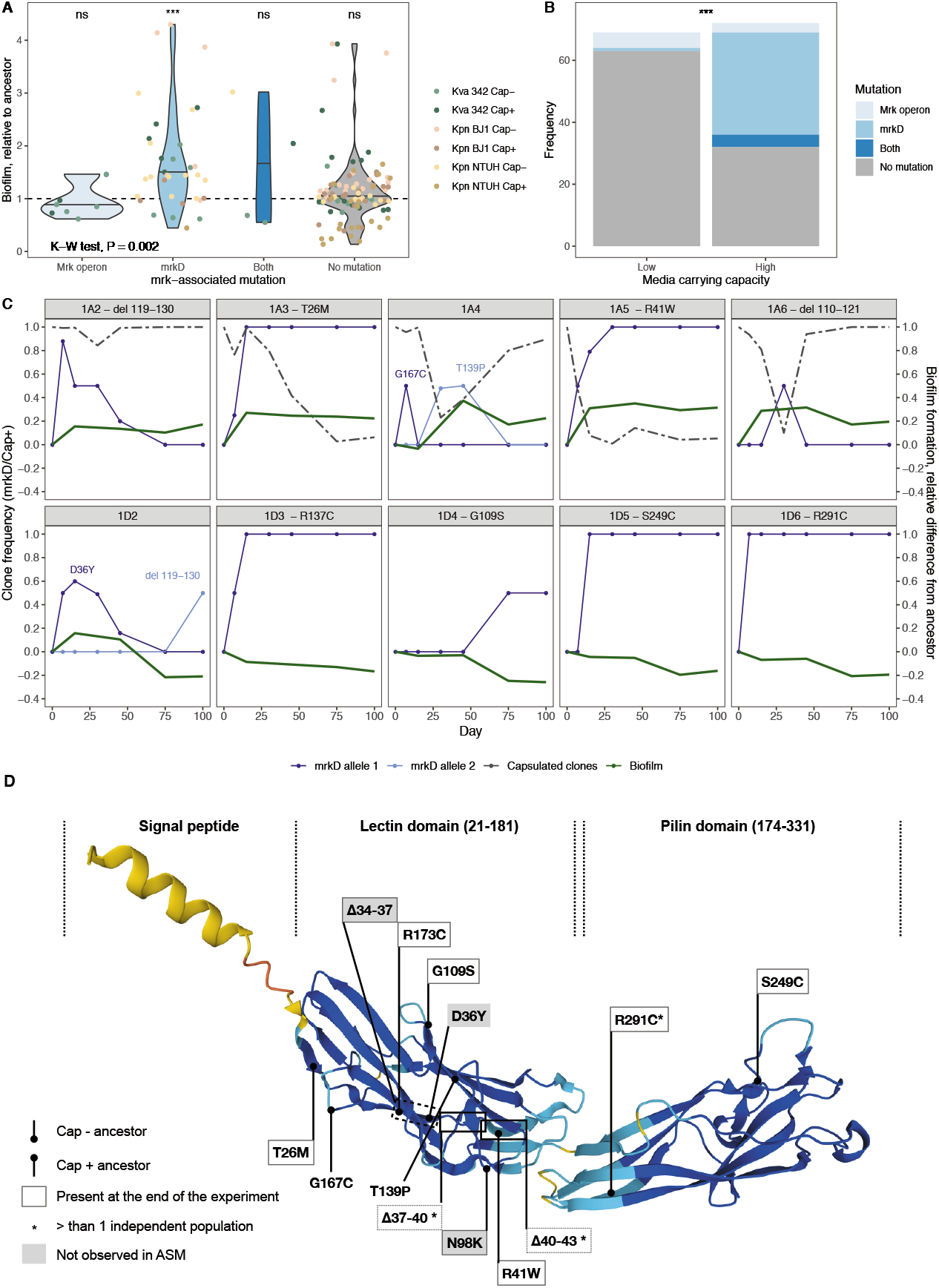
Analyses of evolved populations with mutations in *mrkD*. **A.** Biofilm formation of end-point evolved populations. Populations were divided into four categories corresponding to whether the sequenced randomly-chosen clone from each population had a mutation in *mrkD*, elsewhere in the *mrk* operon, on both *mrkD* and elsewhere, or no mutations in the operon. Each dot represents the average of biofilm formation, relative to its respective ancestor, of at least three independent replicates of each individual evolving population. Line across violin plot indicate the median. Statistics on top of violin plots represent One-sample Wilcoxon Rank Sum test, difference from 1, **P<0.01 and *** P<0.001. **B.** Number of clones in which each category of mutation was found, depending on the carrying capacity of the evolutionary environment: high carrying capacity (nutrient-rich) and low carrying capacity (nutrient-poor). *** P<0.001, Fisher’s Exact test. **C.** Frequency of different *mrkD* alleles in each population as estimated by QSVAnalyzer (blue lines), and capsulated clones (dashed grey lines) as per CFU counts. The difference in biofilm formation between each evolved population and the ancestor is depicted by the green lines. No mutations in *mrkD* were identified in populations 1A1 or 1D1, and are not represented here. **D.** Identified mutations in *mrkD* in Kva 342 strain. Protein structure was predicted with AlphaFold [56]. The pLDDT corresponds to AlphaFold score for confidence per residue. All base-pair deletions resulted in in-frame deletions. Dotted boxes correspond to mutations that were found in multiple populations, but not always observed at the end of the experiment.

The abovementioned correlations were performed based on the sequence of one randomly-chosen clone in the population. To get a finer view of the effect of *mrkD* on biofilm formation, we followed the frequency of *mrkD* alleles in a subset of evolving populations and searched to understand if it correlated with changes in biofilm formation (Figure 3C). We focused on Kva 342, an environmental strain which displayed the highest percentage of populations with mutations in the *mrk* operon (47% vs 22% and 29% in Kpn BJ1 and NTUH respectively) (Table S2, Figure S3), in ASM, a host-mimicking environment. We tested all evolving populations (including those that did not have *mrkD* mutations at the end of the evolution experiment). Surprinsingly, 10 out of the 12 populations had mutations in *mrkD* at some point during the evolution experiments, even if most of them did not reach high frequencies or fixate. Our data revealed that changes in biofilm formation were influenced by the frequency of mutated *mrkD* alleles in the population (GLM, R^2^ = 0.6, P= 0.005).

Because non-capsulated clones readily emerge in capsulated populations [23, 32], we also included the proportion of capsulated clones in the generalized linear model. This revealed that the changes in biofilm formation is also associated with the frequency of capsulated clones (P< 0.0001). We observed that in the two capsulated clones in which the *mrkD* allele fixated (*i.e.* population 1A3 or 1A5), the increase in biofilm formation correlated with an increase in frequency of *mrkD* evolved alleles (Figure 3C). Further, these two populations were the only ones where newly emerged non-capsulated clones outcompeted all capsulated clones by the end of the experiments. In populations 1A4 and 1A6, the emergence of *mrkD* clones correlated with decreased frequency of capsulated clones, despite the fact that neither non-capsulated clones nor *mrkD* alleles fixated. This could suggest that *mrkD* mutations do not emerge in capsulated backgrounds readily. Indeed, among the 168 evolving populations, the mutations in the *mrk* operon are more frequent in the non-capsulated ones (46% vs 19% in capsulated populations, Fisher’s exact test P=0.005). Alternatively, it could suggest that *mrkD* mutations in a capsulated background emerge as readily but do not offer such a large fitness advantage (compared to those in non-capsulated backgrounds) and are outcompeted. To distinguish between the two hypotheses, we isolated six capsulated and six non-capsulated clones from each originally capsulated population at the time point in which the different *mrkD* alleles had reached a frequency of 0.5 in the population (Table S3). Our results show that, *mrkD* mutations emerge both in non-capsulated (populations 1A5 and 1A6) and in capsulated clones (1A2, 1A4 G167C), but they tend to go extinct in the latter. Interestingly, in population 1A3, *mrkD* mutation first emerged in a capsulated background, but our data suggests that shortly afterwards the capsule was inactivated and the non-capsulated clones which encoded a mutation in *mrkD* fixated in the population. Overall, our data shows that mutations in *mrkD* can be associated to changes in biofilm formation, but the latter seems to depend on more complex interactions between the capsule and the different evolved alleles.

### Mutations in *mrkD* increase biofilm formation and reduce aggregation in *K. variicola* but not in *K. pneumoniae*

To disentangle the role of the different *mrkD* alleles in biofilm formation and intercellular interactions, and how the capsule may affect these, we reverted mutations in *mrkD* to the ancestral state in all three strains. The reversion of mutations did not result in consistent changes in biofilm formation in *K. pneumoniae* or in *K. variicola* 342, (Figure S4A and S5A), except for the reversion of Δ37-40 in a capsulated background (1A2) that significantly reduced biofilm formation (Figure S5A). We hypothesized that mutations occurring after the mutation in *mrkD* could mask change in the biofilm phenotype. Indeed, insertion of selected *mrkD* evolved alleles in both capsulated and non-capsulated backgrounds in Kva 342, revealed that *mrkD* mutations increased significantly biofilm formation, but only when the capsule was present (Figure 4A, Figure S5B). This was independent of the genetic context in which the mutation originated, that is, whether it had originally emerged in a capsulated or non-capsulated clone (Multiple-way ANOVA, df = 1, P > 0.05). Given previous studies showing that the capsule limits adhesin exposure [33], we expected mutations in non-capsulated clones to have larger effects on biofilm formation. But, contrary to our expectations, we found no effect (Figure 4A). Analysis of variance confirms that the presence of capsule strongly impacts the effect of *mrkD* mutations (Multiple-way ANOVA, df = 1, P < 0.001). In *K. pneumoniae*, the results were different: only the simultaneous insertion of two SNPs in *mrkD* resulted in significant increases of biofilm formation (Figure S5B). Our data seems to imply that mutations in *mrkD* were not selected for their role in biofilm formation, as the convergent evolution observed in the fixation of these alleles, is not followed by convergent functional evolution.

**Figure 4.**
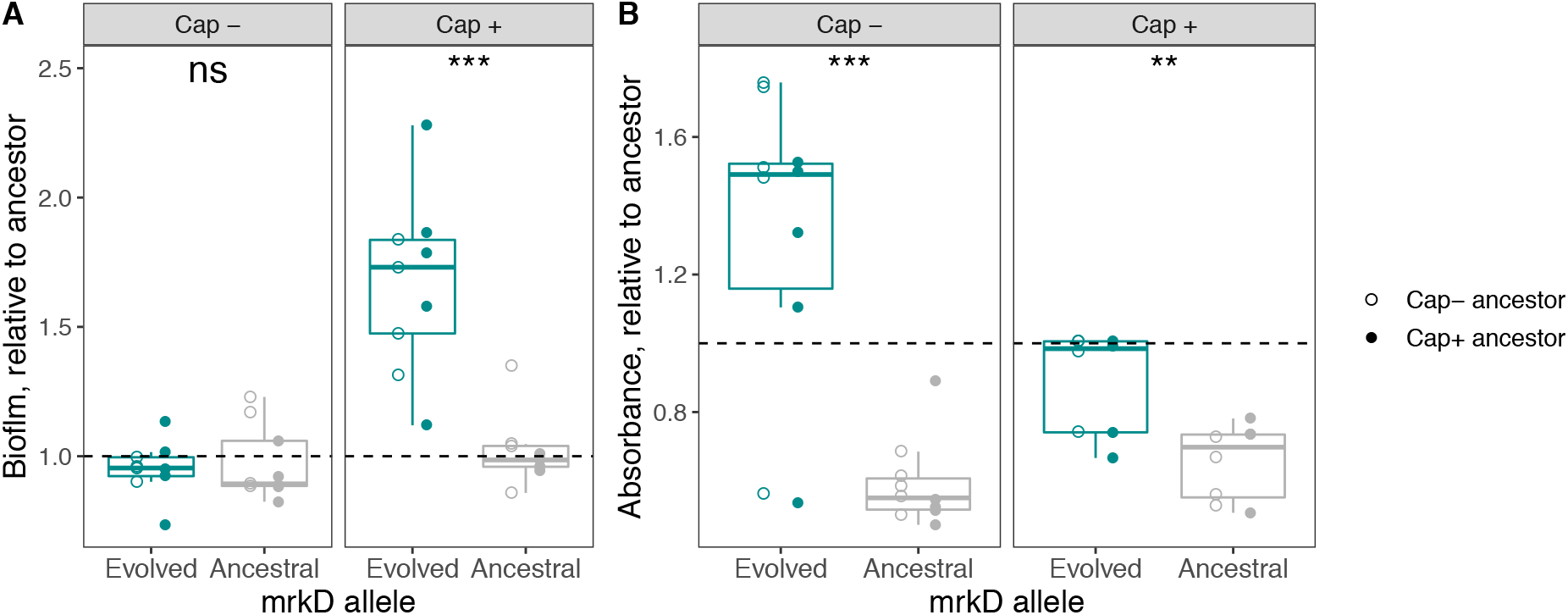
Effect on biofilm formation (A) and aggregation (B) of evolved *mrkD* alleles in ancestral capsulated or non-capsulated Kva 342. Aggregation is quantified by the absorbance (OD600) of the top layer of culture in static conditions after 4.5 hours. High absorbance results from low aggregation levels. Calculation of the area under the aggregation curve results in qualitatively similar results. Each dot represents a clone with an independently evolved mutation. Full points represent mutants in which *mrkD* allele originally emerged in a clone descending from a capsulated ancestor whereas empty points represent mutants bearing *mrkD* alleles that emerged in non-capsulated ancestors. For individual visualization of each mutation, and individual statistics see Figure S5. Statistics: two-sided paired t-tests. *P<0.05,**P<0.01, ***P<0.001

Because most mutations in *mrkD* were found in the lectin (collagen) binding domain (Figure 3D), we tested whether these mutations could specifically impact cell-to cell interactions. To do so, we measured the absorbance of the top layer of sitting cultures through time as a proxy for sedimentation. In such tests, high absorbance represents low sedimentation. Aggregation results confirm what was observed in biofilm formation, namely that in Kva 342, the *mrkD* allele (Multiple-way ANOVA, df = 1, P < 0.001), as well as the presence of the capsule (Multiple-way ANOVA, df = 1, P = 0.008), but not the ancestral background in which the mutation originally emerged (Multiple-way ANOVA, df = 1, P > 0.05), impacts aggregation. Specifically, most mutations significantly reduced aggregation relative to the ancestor in Kva342 (Figure 4B, Figure S5C). No changes in aggregation were observed in *mrkD* mutants of *K. pneumoniae*.

Taken together, our data show that irrespective of the ancestral capsule genotype, mutations in *mrkD* increase biofilm formation in capsulated clones whilst diminishing cell-to-cell interactions in *K. variicola* but not in *K. pneumoniae*.

### Non-capsulated *K. pneumoniae* populations revert the *fim* switch and form less biofilm

Analyses of mutations observed in end-point populations revealed that some Kpn NTUH population had clones that reversed the *fim* switch. The *fim* switch is a phase-variable inversion of a short DNA element (comprising the promoter), which results in an ON/OFF expression of the *fim* operon, responsible for the production, or not, of type I fimbriae [34]. Seventeen out of the 56 Kpn NTUH evolved populations had clones with reverted *fim* switch and these were predominant in the population. The reversions of the promoter did not depend on the ancestral capsule genotype, but were dependent on the environment in which the population evolved. Switch reversal accumulated preferentially in environments with low-carrying capacities (Fisher’s test, P=0.002), as opposed to mutations in *mrkD* which mostly accumulated in environments with high carrying-capacities (Figure 5)

**Figure 5.**
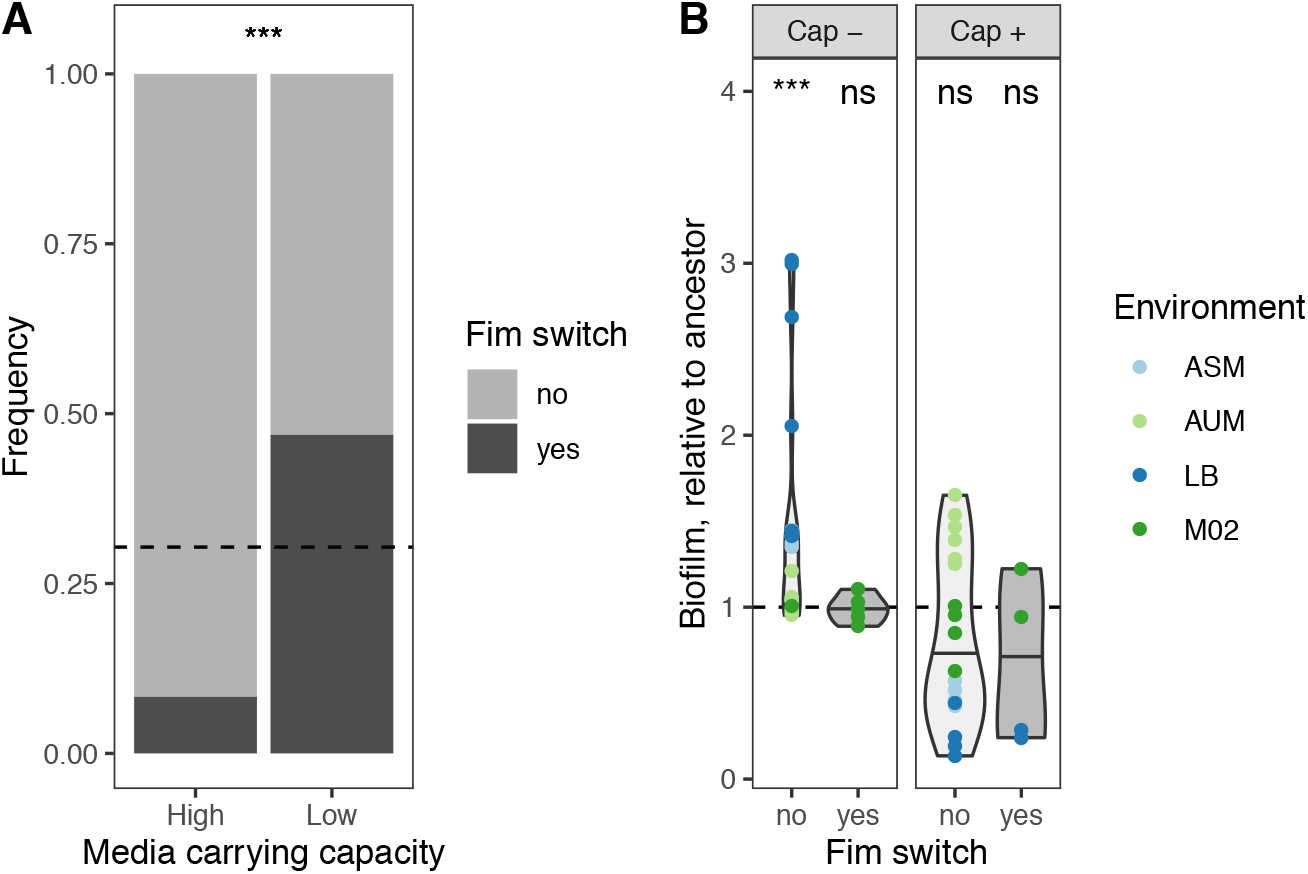
Analyses of reversal of *fim* switch in evolved populations. **(A)** Tropism of *fim* switch reversal. The dashed horizontal line indicates the total frequency of *fim* switch in evolved clones. Fishers Exact test was used to compute difference in the distribution of mutations depending on media carrying capacity. ***P<0.001 **(B)** Biofilm formation was assessed in the evolutionary treatment in which this mutation evolved. Each dot represents the average of three independent experiments of each individual population. Blue dots depict populations evolved in media with high carrying capacity and green indicates media with low carrying capacity. Statistics: Two-tailed One-sample t-test, difference from 1. ***P<0.001, ns: not significant.

We then tested whether Kpn NTUH populations with the reversed *fim* promoter displayed changes in biofilm formation. Analysis of variance revealed that variation in biofilm formation was not dependent on the environment (Multi-way ANOVA, df = 3, P> 0.05) nor on mutations in the *fim* promoter (Multi-way ANOVA, df = 1 and P> 0.05). In contrast, it was strongly associated with the ancestral capsule genotype (Multiway ANOVA, df = 1, P = 0.0003). Whereas in capsulated backgrounds, populations displaying changes in the *fim* switch did not result in differences in biofilm formation, in non-capsulated populations, populations with switch reversal produced less biofilm than those without (Figure 5B). Taken together, despite the fact that the reversion of the *fim* switch is equally frequent in capsulated and in non-capsulated populations, it only seems to reduce biofilm formation in non-capsulated *K. pneumoniae* NTUH.

### Mutations in *mrkD* are also found in natural and clinical isolates

The frequency at which *mrkD* mutations were observed prompted us to enquire whether mutations driving latent phenotypes in laboratory evolution experiments could reflect evolutionary paths occurring in natural isolates. To do so, we analyzed all *K. variicola* and almost 10 000 random *K. pneumoniae* genomes available in the Pathosystems Resource Integration Center (PATRIC) genome database [35]. In *K. variicola*, we identified a total 689 MrkD proteins in 671 genomes (prevalence of 93.6%) (Figure S6). From these, 384 protein sequences differed from our ancestral Kva 342. A total of 55 different aminoacid changes were identified, grouped in 42 unique MrkD proteic sequences (Table 1). Among these, three mutations (found in ten different genomes) displayed aminoacid changes in the same positions as proteins evolved in our evolution experiment (position #57, #73 and #249) (Figure 6, Table 1). Genomes with similar mutations were all host-associated, isolated from humans or cats. Further, one human sample came from lung sputum, suggesting that our ASM environment mimics natural conditions. In *K. pneumoniae*, we found a similarly high prevalence of MrkD (93.7%). Fourteen out of the 22 (∼63%) different aminoacid positions that were mutated in our evolution experiment, were also mutated in the natural isolates (Figure 6, Table 1). Taken together, the comparison of the mutations in MrkD in wild isolates and that observed in the laboratory reveals that the evolution experiments captured a broad range of the genetic diversity observed in natural populations.

**Table 1.** Summary statistics of genomes and comparison of MrkD sequences analysed from the public repository PATRIC.

| Species | # of genomes | # mrkD sequences (evalue < 10e<sup>-5</sup>) | Prevalence (> 90% identity) | # sequences <100 %>90% identity | # Unique alleles | #common positions modified | Positions |
| --- | --- | --- | --- | --- | --- | --- | --- |
| *K. variicola* | 717 | 1661 | 671 | 384 | 42 | 3 | 57,73, 249 |
| *K. pneumoniae* | 9761 | 26065 | 9154 | 6794 | 237 | 14 | 26, 36, 37, 41, 57, 58,70, 73, 98, 109, 137, 166, 185, 249 |

**Figure 6.**
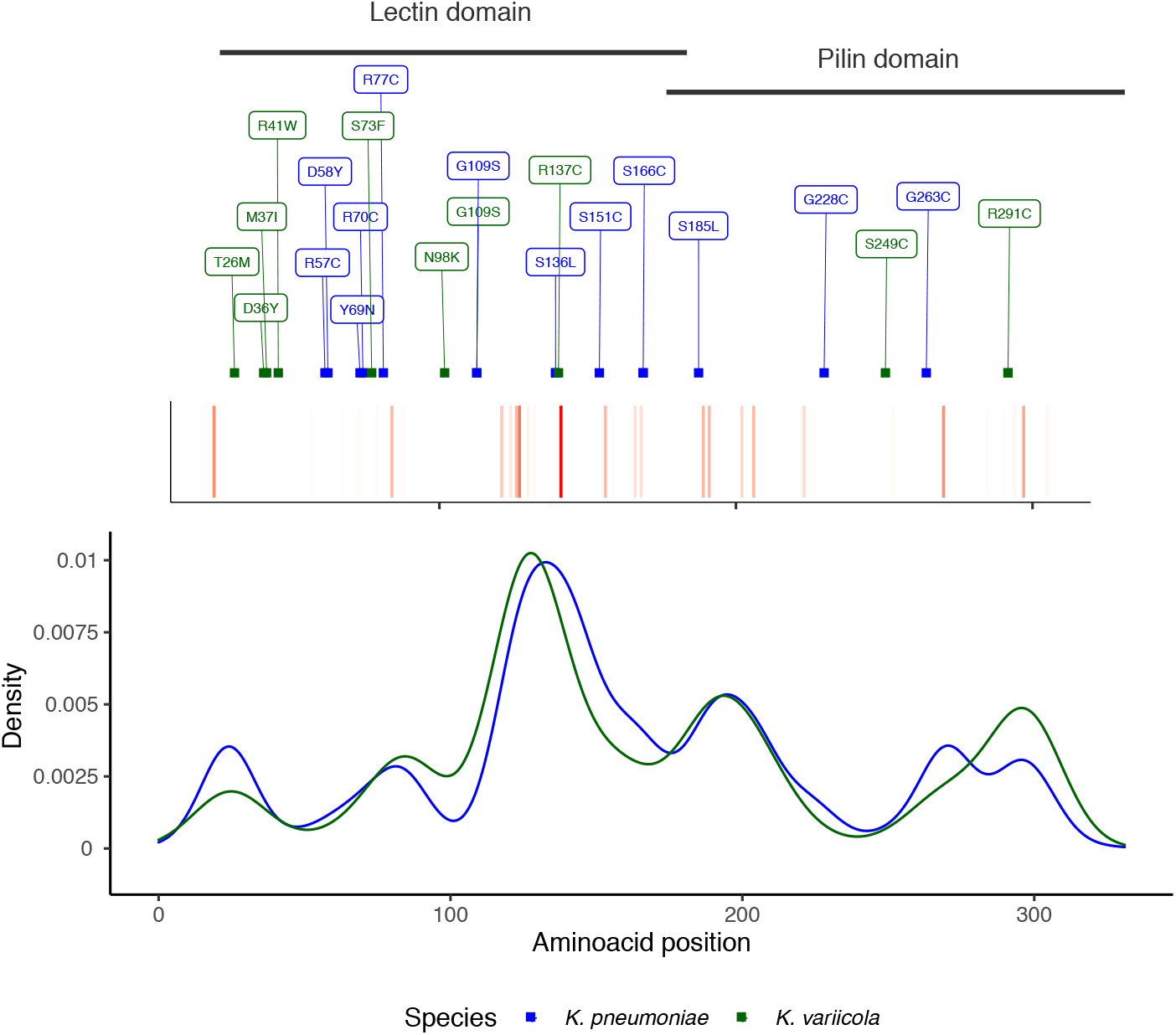
Comparison of MrkD mutations observed in natural isolates and our evolution experiment. Smoothed density plot (default parameters) of the distribution of SNPs in MrkD from *K. variicola* (green) and *K. pneumoniae* (blue). The inset recapitulates the frequency of aminoacid changes in each position. The intensity of red represents the frequency.

## DISCUSSION

Latent phenotypes evolve neutrally during adaptation to novel environments. Such phenotypes do not significantly impact fitness, but they can be adaptive if the environmental conditions change. Recently, numerous experimental evolution studies have tested how adaptation to novel environments may impact latent phenotypes as they contribute to diversification [36, 37] or to exaptation [38, 39]. Indeed, the resulting genetic variation has been called “evolution’s hidden substrate” [40]. Here we studied the diversification and evolution of one such latent phenotype, namely the ability to form biofilm. To do so, we examined 144 evolving populations, grown in static liquid cultures for over 675 generations. Prior to daily passaging, cultures were vigorously homogenized, and thus biofilm formation either at the bottom of the well or in the air-liquid interface was not selected for. We evaluated how the environment and ancestral genotypes impact changes in the trait and how the latter coevolves with other traits. We observed little within-treatment diversity in the evolutionary outcomes in latent evolution of biofilm formation. This contrasted with the divergent evolutionary outcomes observed across treatments. These were driven by the ancestral genotypes, but more importantly by nutrient availability.

The presence or absence of the capsule not only impacts the magnitude of change of adaptive phenotypes like yield at the end of the experiment, as shown previously [32]. It also impacts the evolutionary dynamics of latent phenotypes. The effects of the capsule on biofilm formation seem to be stronger at longer evolutionary time scales. For example, in the first one hundred generations, both capsulated and non-capsulated populations across all environments followed a similar dynamic of fast and large changes in biofilm formation. The impact of the capsule in biofilm dynamics could be a consequence of its effect on an adaptive trait, like increased population yield. Changes in yield are frequent at the beginning of evolution experiments [41]. This is expected as the most beneficial mutations are incorporated early in the process [42]. At longer evolutionary time scales, divergent evolutionary paths are observed in terms of biofilm formation depending on the presence of the capsule and on the carrying capacity of the environment. This suggests that these two factors critically alter the future ability of microbes to form biofilm, which affects the way they prevail and better colonize a surface (including the host).

We hypothesized that the convergent accumulation of mutations across the three strains in the type 3 fimbriae tip adhesin *mrkD*, would phenotypically impact the formation of biofilm because MrkD mediates the adhesion to cells and several extracellular matrix proteins [43]. However, the phenotypic consequences of such mutations were not similar. Specifically, we observed that *mrkD* mutations increased biofilm formation in the *K. variicola* strain, but not in the *K. pneumoniae* strains. Further, we observed that reversions of *fim* switch accumulate in only one *K. pneumoniae* (NTUH), but not in BJ1 or in the *K. variicola* strain. This suggests that the importance of this adhesin in biofilm formation could be strain-dependent. Computational analyses have revealed that *K. pneumoniae* strains have a rich arsenal of chaperon-usher (CU) systems. Most of these systems are cryptic in laboratory conditions [21] and little is known about their expression and conservation patterns. In light of our results, further research evaluating the precise role of each system in the presence and absence of the other CU-systems would be a promising research venue to unravel the inter-adhesin interactions and how they influence biofilm formation. The relative expression of each CU-system across different strains and environments could significantly alter the contribution of each system in biofilm formation [21].

A major finding of this study is that differences in nutrient availability led to the differential accumulation of mutations in *mrkD*. This raises the question of why *mrkD* mutations are selected specifically in nutrient rich but not in nutrient poor environments? We speculate that in nutrient rich medium, when higher carrying capacities are sustained, fitness advantages may often be the result of higher growth rates. Because single cells have higher intrinsic growth rates than aggregates, mutations leading to decreased cell aggregation would be selected [44]. However, when competition is high and resources scarce, cell aggregates have higher fitness than single cells [44], thus mutations in *mrkD* may be counter-selected in nutrient-poor environments. Alternatively, there may be a specific tropism of the fimbriae. Selection can only act on what is being expressed in each environment. Thus, we can speculate that changes in fimbriae expression across the different environments are driving mechanisms of adaptation. For instance, we observe reversion of the *fim* switch in nutrient-poor environments, like AUM, where they are expressed and known to play an important role [45]. On the contrary, we do not observe many reversion of the *fim* switch in nutrient-rich environments, where they are not expressed *i.e.* during lung infection [45]. A similar tropism could explain accumulation of mutations in *mrkD* in certain environments but not in others. Finally, given that the mutations in *mrkD* do not result in consistent changes in biofilm formation and/or aggregation across ancestral genotypes (strain and capsule), we may also speculate that selection is acting on another phenotype influenced by *mrkD* and of particular relevance in nutrient-rich environments. If so, it would suggest that this CU-system may fulfil roles unrelated to biofilm formation.

Our study reveals an intricate relationship between the capsule and extracellular proteins. The effect of the *fim* switch is only visible in non-capsulated cells. This is not unexpected as it was initially shown that the capsule in *Klebsiella* limited biofilm formation, notably by masking the effect of type I fimbriae [33] and other short adhesins such as Ag-43 and AIDA-1[24]. Accordingly, when the capsule contributes to biofilms’ maturation and is highly expressed [22], type I fimbriae are downregulated [16] and not involved in intestine or lung colonization [34]. However, in these conditions, type 3 fimbriae are highly expressed, suggesting a positive interaction between the capsule and this fimbriae during biofilm formation. Similarly, a recent report showed that, in conditions in which type 3 fimbriae is expressed, biofilm formation increases when capsule production is enhanced [46]. Indeed, in *K. variicola*, the mutations in MrkD only significantly increase biofilm in capsulated strains, but not in non-capsulated strains. Such differential increase of biofilm is interesting, as *mrkD* mutations indiscriminately impact aggregation irrespective of the presence or absence of the capsule. This highlights that the interaction between the capsule and type 3 fimbriae during biofilm formation cannot be reduced to a mere masking of surface receptors and depending on the context, they may act synergistically.

The convergent evolution of mutations on the lectin domain of the tip adhesin MrkD is similar to that recently observed in the lectin domain of the tip adhesin FimH of type I fimbriae in *E. coli*, in a study in which biofilm formation under continuous flow was positively selected [30]. The nature of the mutations was also found to be very similar between the two studies, especially the emergence of several short in-frame deletions across independent populations. Of note, some clones with mutations in *fimH*, as observed with clones with mutations in *mrkD*, did not significantly increase biofilm formation relative to the ancestor. This reinforces our hypothesis that there may be other selective pressures underlying the emergence and rise to such high proportions of mutations in the tip adhesin of chaperon-usher dependent fimbriae. Alternatively, they may play roles other than formation of biofilm, either directly or as a result of epistasis with other surface structures. Despite the differences in selection regimes between the two studies, the mutagenic convergence in the lectin domain of the tip adhesins in *Klebsiella* and *E. coli* suggests that proteins with similar functions undergo similar evolutionary trajectories.

At the population level, we observed a remarkable correlation between the presence of mutations in the tip adhesin *mrkD* and increased biofilm formation (Figure 3A). Yet, this effect could not be fully recapitulated when these mutations were analysed individually in an ancestral background. This could be due to complex epistatic interactions, for instance, with the capsule or other surface structures. Alternatively, the effects of these mutations could be altered by the presence of other mutations in the genome. We had previously reported that some populations also had mutations in *ramA* [32], involved in the stability of the outer membrane [47]. These could alter molecular interactions at the cell surface, including the adhesin-capsule interactions. Further, in *Salmonella enterica* Typhimurium, expression of *ramA* directly impacts biofilm formation [48]. Finally, the formation of biofilm can be regarded as a social behaviour and thus, strongly influenced by the within-population diversity. Indeed, here we analysed genotypes in isolation, but clonal interference may be influencing changes observed at the population level.

The impact of direct selection on important virulence-associated traits has been largely studied, yet, changes in traits that evolve latently are rarely addressed. This could be caused by the challenges of analysing the genetic basis of their evolution, since these phenotypes may not have a direct impact on fitness. The phenotypic changes resulting from specific mutations may not be easy to reveal and may strongly rely on complex epistatic interactions or within-population diversity. Previously, we showed how susceptibility to antimicrobials latently evolves as a result of mutations in *ramA* [32]. Here, we have shown that the ability to form biofilm changes greatly during evolutionary time, depending on the ecological conditions in which *mrkD* mutations emerge, and on the genetic context in which it is expressed. In conclusion, our studies highlight the need to include analyses of latent phenotypic evolution in the general framework of microbial adaptation, and more particularly in the evolution of virulence-associated traits of bacterial pathogens [36, 37].

## MATERIALS & METHODS

### Bacterial strains and growth conditions

*i. Strain*. Three different strains from the *Klebsiella pneumoniae* species complex were used in this study [32]: one environmental strain, isolated from maize in the USA, *K. variicola* 342 (Kva 342, serotype K30) [47], *K. pneumoniae* BJ1 from clonal group 380 (Kpn BJ1, serotype K2) isolated in France from a liver abscess [11] and the hypervirulent *K. pneumoniae* NTUH K2044 (Kpn NTUH, serotype K1) from clonal group 23 isolated in Taiwan from a liver abscess [48]. *ii. Environment description.* AUM (artificial urine medium) and ASM (artificial sputum medium) were prepared as described previously [49, 50]. AUM is mainly composed of 1% urea and 0.1% peptone with trace amounts of lactic acid, uric acid, creatinine and peptone. ASM is composed of 0.5% mucin, 0.4% DNA, 0.5% egg yolk and 0.2% aminoacids. LB is composed of 1% tryptone, 1% NaCl and 0.5% yeast extract. M02 corresponds to minimal M63B1 supplemented with 0.2% of glucose as the sole carbon source. *iii. Primers.* Primers used in this study are listed in Table S4.

### Mutant construction

Isogenic mutants were constructed by allelic exchange. We inserted evolved *mrkD* alleles in capsulated and non-capsulated ancestors. We also reverted evolved alleles into ancestral state in evolved clones. To do so, the cloning vector pKNG101 plasmid was amplified using Phusion Master Mix (Thermo Scientific) and digested by DpnI (NEB BioEngland) restriction enzyme for 30 minutes at 37°C. The *mrkD* allele of interest (ancestor or evolved) was amplified using Phusion Master Mix (Thermo Scientific). The vector and allele of interest were then assembled using the GeneArt™ Gibson Assembly HiFi kit (Invitrogen), electroporated into competent *E. coli* DH5α strain and selected on streptomycin LB plates. pKNG101 plasmids containing *mrkD* alleles were verified by PCR and purified using the QIAprep Spin Miniprep Kit. These were then electroporated again into *E. coli* MFD λ-pir strain, used as a donor strain for conjugation in Kva 342 ancestral strains or evolved clones. Single cross-over mutants (transconjugants) were selected on Streptomycin plates and double cross- over mutants were selected on LB without salt, supplemented with 5% sucrose, after 48h of growth at room temperature. From each double-recombination, an evolved alelle and an ancestral allele were isolated. Mutants were verified by Sanger sequencing.

### Emergence of *mrk* mutants

To test the allele frequencies of *mrkD* in the evolved populations, we took aliquots from the glycerol stock from days 7, 15, 30, 45, 75 and 100. These were diluted in water and used for PCR reaction using Phusion Master Mix (Thermo Scientific). Purified PCR product was sent for Sanger sequencing. The frequency of the mutations was calculated using high quality chromatograms analyzed by QSVanalyzer [51] with default parameters. Of note, the limit of detection of QSVanalyzer is estimated at 5%.

### Trait quantification

To initiate the different measurements, each population was grown overnight and 20 μL of each culture was inoculated into 1980 μL of the relevant growth media in 24-well microtiter plates and allowed to grow for 24 hours without shaking at 37°C. Populations evolving in poor media (M02 and AUM) were diluted 1:100 and allowed to grow for another 24-hours extra prior to the experiment, as we noticed that preconditioning was important for reproducibility. (i)*Biofilm formation*. The capacity of a population or isolated clones to form a biofilm was performed as previously described [52], with volume modification. Briefly, unbound cells were removed by washing once in distilled water. To stain biofilms, 2100 μL of 1% crystal violet was added to each well for 20 minutes. The crystal violet was decanted and washed thrice with distilled water. The plates were allowed to dry under a laminar flow hood. Then, the biofilm was solubilized for 10 min in 2300 μL of mix with 80% ethanol and 20% acetone. Two hundred μL of each mix was transferred in a well of a 96-well plate.

The absorbance of the sample was read at OD_590nm_. (ii)*Population yield*. Each well was homogenized by vigorous pipetting and then serially diluted in fresh LB and plated to count CFU after 24 hours of growth. For most capsulated populations in LB and ASM, due to the extreme hypermucoviscous phenotype [32], CFU could not be accurately assessed as the populations cannot be resuspended and homogenized. Thus, the serial dilution process was biased and distorted because either a randomly large or a randomly small proportion of the population would be transferred, due to the abovementioned hypermucoviscosity. These populations were not taken into account in our analyses (iii) *Surface polysaccharide extraction and quantification.* The bacterial capsule was extracted as described in [53] and quantified by the uronic acid method [54] using glucuronic acid as a standard. (iv) *Aggregation test.* An isolated colony was allowed to grow in 5 mL overnight in M02 medium at 37° under shaking conditions. Prior to the experiment, the absorbance (OD_600nm_) was measured and adjusted to OD_600_ = 2, and the cultures were transferred to static test tubes. Two hundred μL samples were transferred to a 96-well microtiter plate and the absorbance (OD_600nm_) measured at defined time points (0 ; 1,5 ; 3 ; 4,5 and 24 hours) using an automatic plate reader Spark Control Magellan (TECAN). Samples were removed from the uppermost layer of tube cultures, roughly at the 4 mL mark. Decreasing absorbance represents the settling of agglutinated cell clumps. Figures represent aggregation after 4.5h. We also calculated the area under the aggregation curve. Results were qualitatively similar to those observed when only the measurement of 4.5h is taken into account.

### Search for MrkD proteins

All genomes corresponding to *K. variicola* in the Pathosystems Resource Integration Center (PATRIC) genome database, filtered by good quality and text mined for *K. variicola* species (751 out of the 767) were downloaded on March 11, 2022. Same procedure was applied for the first 10 000 genomes of *K. pneumoniae* (of which 239 were discarded). The genomes were checked for quality control and annotated with the pipeline PaNaCoTa [55] and the --prodigal option. Protein-Protein Blast (BLAST 2.7.1+) against either the Kva 342 MrkD or Kpn NTUH MrkD was performed with the following option - max_target_seqs 100000 and an E-value smaller than 10^-5^. Sequences covering less than 70% of the protein length were discarded (only 50 for *K. pneumoniae*). The distribution of MrkD protein identities revealed a bimodal distribution (Figure S6A, C). To analyse *bona fide* MrkD proteins, we applied a cut-off of 90% identity. Most of the unique MrkD protein sequences in the databases differed by one or two aminoacids from our ancestral sequences, but these could go up to 20 (Figure S6B, D). Proteic sequences were aligned using mafft v7.22 with the options --localpair --maxiterate 1000. Aminoacid mismatches were then identified using the R package Biostrings.

## DATA ACCESIBILITY

Raw data is available in the figshare repository under the doi 10.6084/m9.figshare.22559890. Raw reads for this project can be accessed in the European Nucleotide Archive (ENA), project number PRJEB54810 [32].

## SUPPLEMENTAL FIGURES

**Figure S1.**
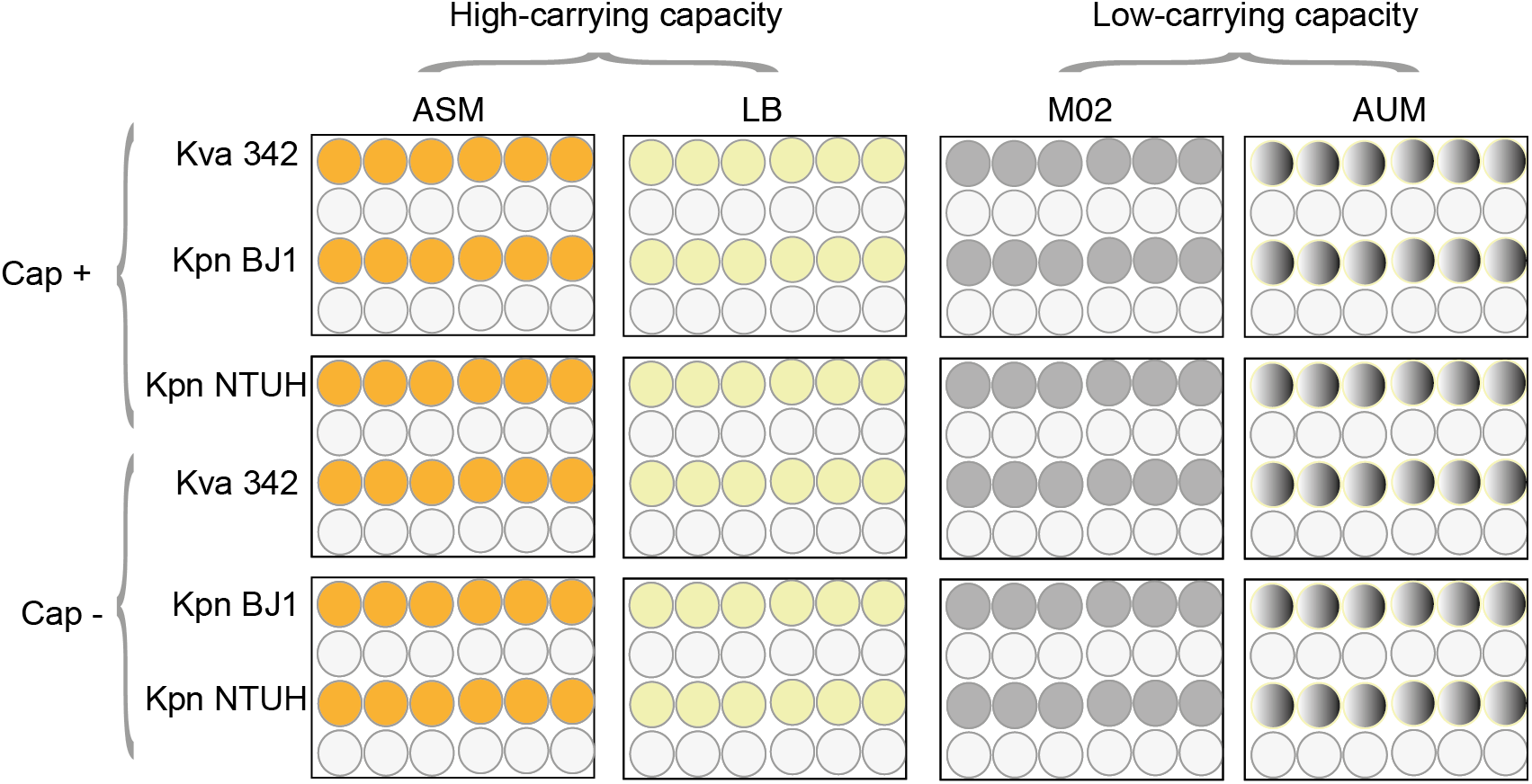
Schematic representation of the evolution experiment. Six replicate populations of each genotype were serially passaged over 102 days. After 24 hours at 37° in static conditions, vigorous homogenisation by pipetting was applied to each population and 1% (20μL) was transferred into a new well containing 1980 μL of fresh media. One empty row was left between strains to avoid cross-contaminations. For the purpose of this study, the soil environment included in the original evolution experiment [32]was not examined as biofilm formation in soil is below the limit of detection of the assay.

**Figure S2.**
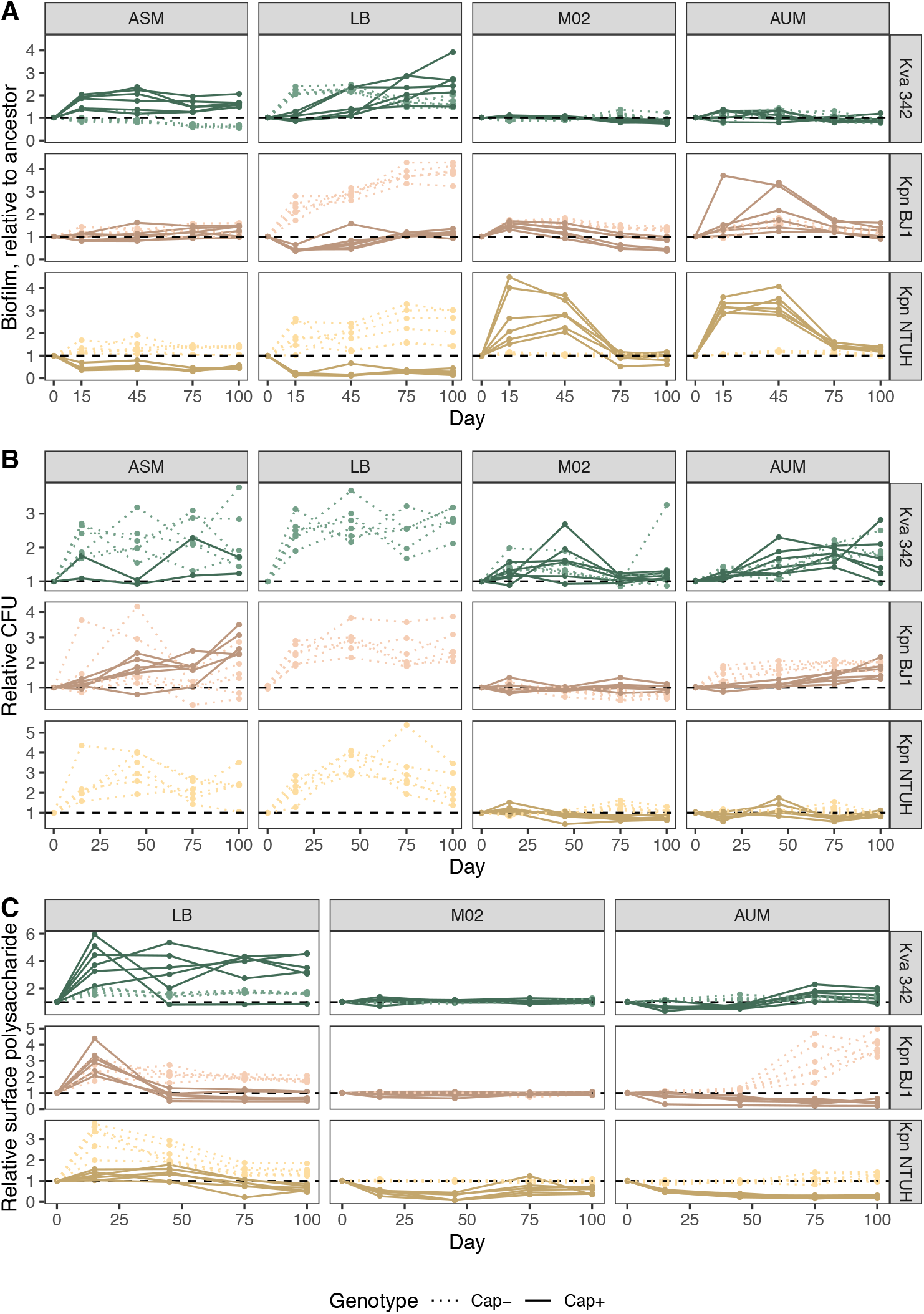
Changes in biofilm formation (A), population yield (B) and production of surface polysaccharides (C), relative to the ancestor. Biofilm formation, population yield and surface polysaccharide production of each individual evolving population was tested, relative to its ancestor. Measures were taken at day 15,45, 75 and 102. Each point is the average of at least three independent biological replicates. Error bars are not shown for visualization purposes. Population yield could not be assessed in some capsulated populations in high- carrying capacity environments (ASM & LB) because populations were hypermucoviscous, precluding any reliable assessment [32]. Surface polysaccharides in ASM could not be measured as some components of ASM interfered with the measurements. However, the presence of such components varied across time and populations (probably due to bacterial consumption) and thus surface polysaccharides could not be compared across populations.

**Figure S3.**
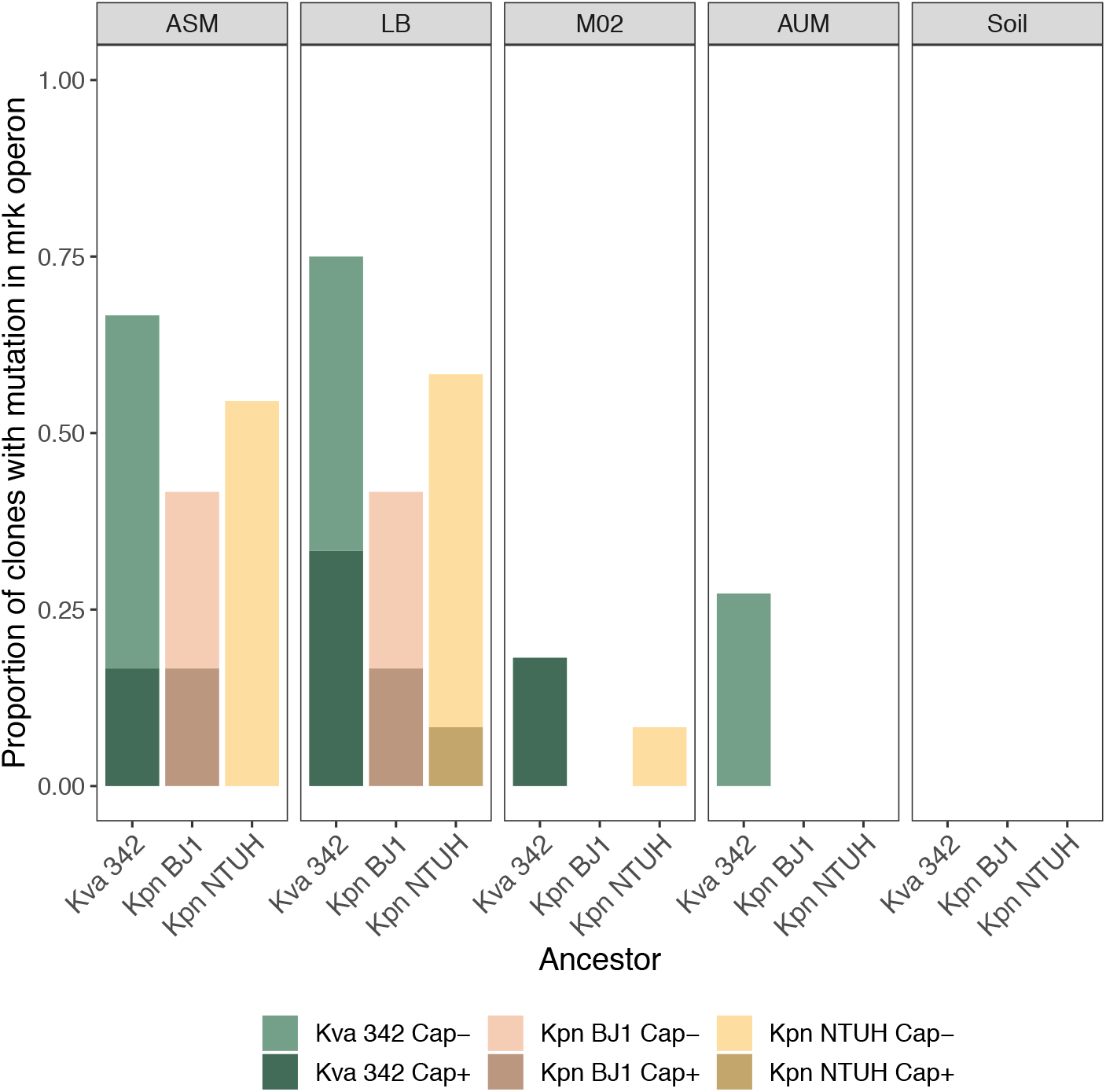
Proportion of individual clones with mutations in the *mrk* operon across ancestors and environments.

**Figure S4.**
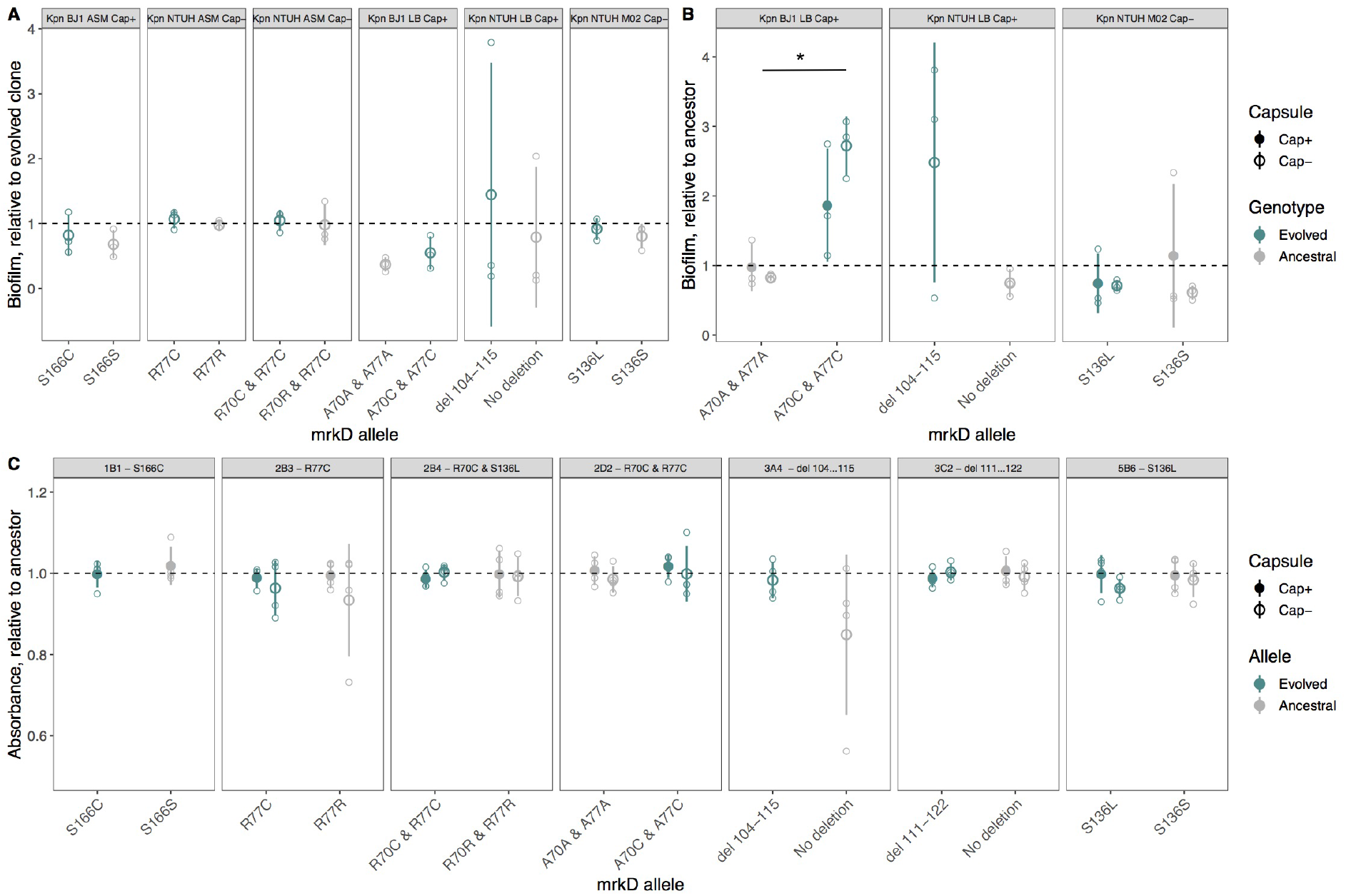
Effect of *mrkD* mutations in *K. pneumoniae* BJ1 and NTUH on biofilm formation (A and B) and aggregation (C). Biofilm formation was assessed in the evolutionary treatment in which these mutations emerged (ASM, LB or M02). Full large points represent the average of capsulated clones and empty points represent non-capsulated clones. Grey points represent ancestral allele whereas green points indicate evolved alleles. Statistical analysis was performed to compare evolved vs ancestral alleles, two-sided paired t-test. Only significant comparisons are indicated * P<0.05.

**Figure S5.**
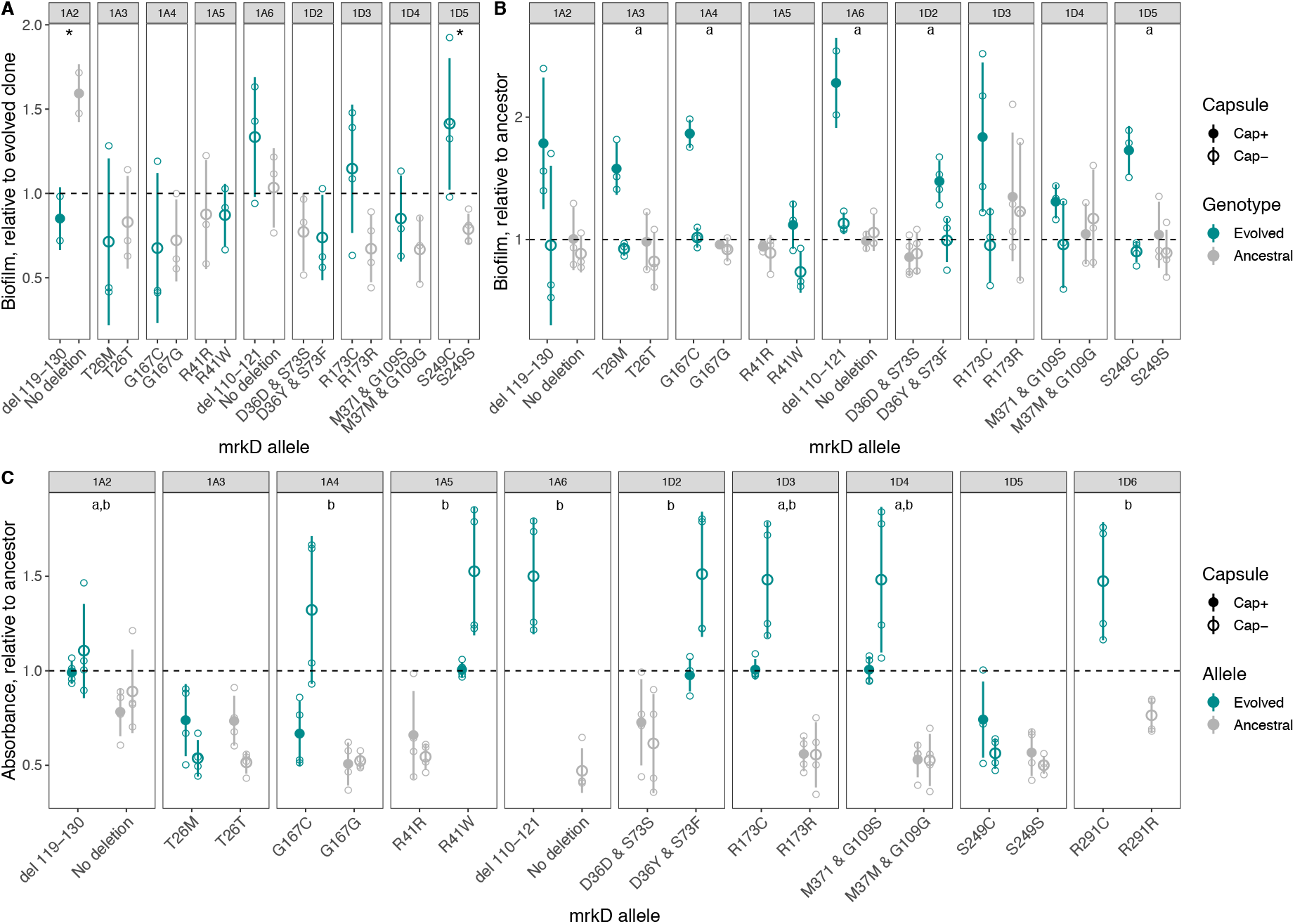
Effect of *mrkD* mutations in *K. variicola* 342 on biofilm formation (A and B), aggregation (C). Panels A represents reversion of evolved allele into ancestral allele in the evolved clones. Of note, some reverted clones from originally capsulated populations are non-capsulated (1A3,1A4,1A5 and 1A6). Panels B and C represent insertion of evolved allele in ancestral backgrounds (capsulated and non-capsulated backgrounds). Full large points represent the average of capsulated clones and empty point represent non-capsulated points. Grey points represent ancestral allele whereas green indicate evolved alleles. Small individual points indicate independent experiments. Statistical analysis was performed to compare evolved vs ancestral alleles, two-sided paired T-test. ‘a’ indicates P<0.05 across capsulated clones whereas ‘b’ indicates P<0.05 across non-capsulated clones. When only one comparison was necessary * P<0.05, **P<0.01. Only significant comparisons are indicated.

**Figure S6.**
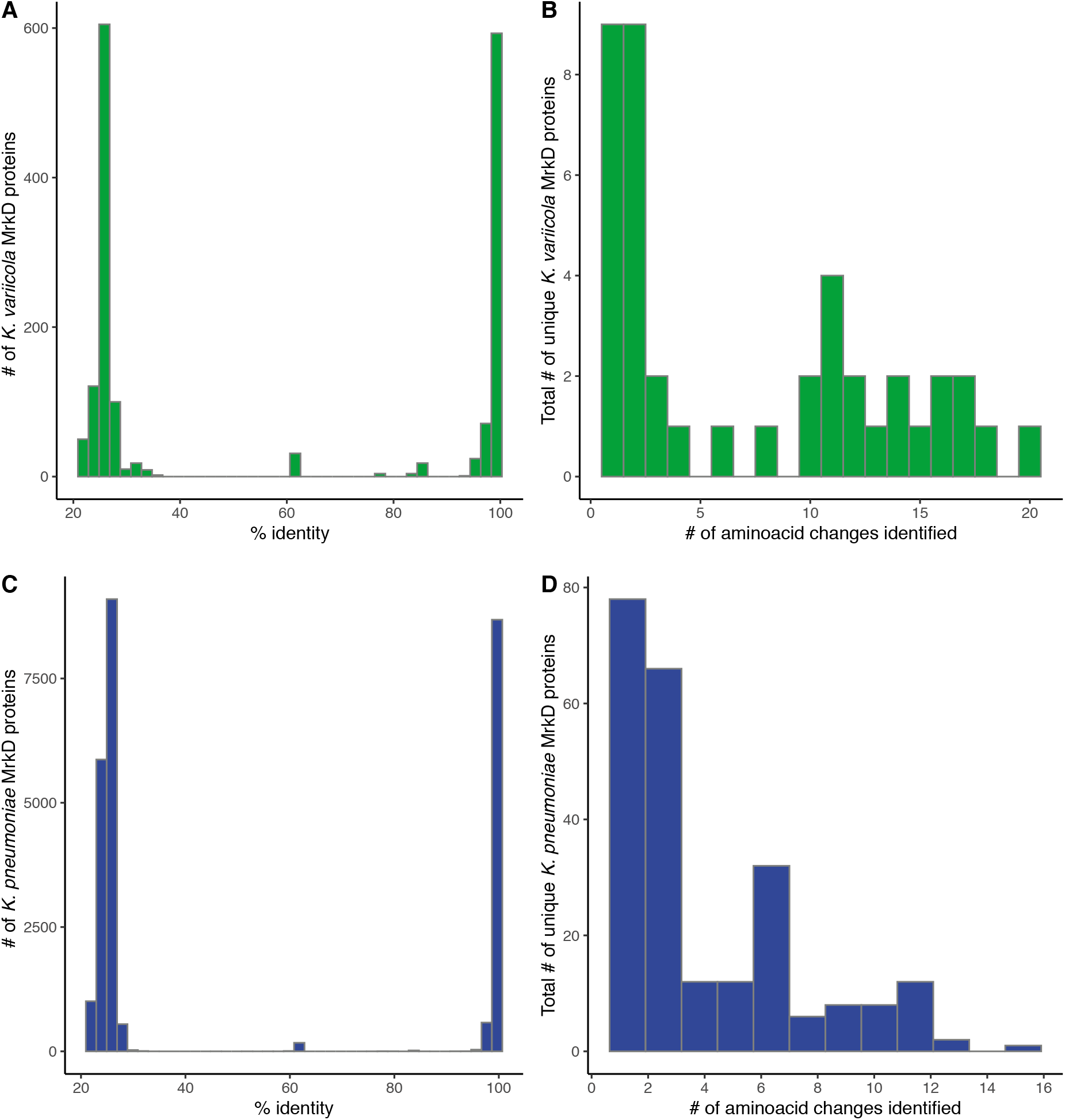
Diversity of *mrkD* alleles in genomic databases. Distribution of the protein sequence identity between all proteins found in the databases for *K. variicola* (**A**) and *K. pneumoniae* (**C**). Proteins were identified by pblast with an e-value of less that 10^-5^. Only proteins with more than 90% identity were selected for further study. The total number of aminoacid differences between our ancestral sequences and those found in the databases are depicted for *K. variicola* (**B**) and the *K. pneumoniae* (**D**).

## SUPPLEMENTAL TABLES

**Table S1.** Changes in population yield and surface polysaccharide production relative to the ancestor. Statistics correspond to One-sample Wilcoxon Rank Sum test. CI95 indicates the interval of confidence. Highlighted in italics are those traits that did not significantly increase, relative to the ancestor.

| Day | Trait | Carrying capacity | N | Median | Mean | CI95 | P-value |
| --- | --- | --- | --- | --- | --- | --- | --- |
| 15 | Relative CFU | Low | 72 | 1.13 | 1.15 | 0.13 | <0.001 |
| 15 | Relative CFU | High | 44 | 1.96 | 2 | 1.06 | <0.001 |
| 15 | Relative surf. polysacch | Low | 72 | 0.9 | 0.87 | 1.03 | 0.001 |
| 15 | Relative surf. polysacch | High | 36 | 2.55 | 2.63 | 2.07 | <0.001 |
| 45 | Relative CFU | Low | 72 | 1.17 | 1.23 | 1.34 | <0.001 |
| 45 | Relative CFU | High | 44 | 2.54 | 2.54 | 1.62 | <0.001 |
| 45 | Relative surf. polysacch | Low | 72 | 0.9 | 0.85 | 1.57 | 0.003 |
| 45 | Relative surf. polysacch | High | 36 | 1.77 | 1.9 | 1.22 | <0.001 |
| 75 | Relative CFU | Low | 72 | 1.22 | 1.24 | 0.55 | <0.001 |
| 75 | Relative CFU | High | 44 | 2.11 | 2.14 | 1.09 | <0.001 |
| 75 | *Relative surf. polysacch* | *Low* | 72 | *1* | *1.1* | *1.06* | *0.882* |
| 75 | Relative surf. polysacch | High | 36 | 1.53 | 1.7 | 0.76 | <0.001 |
| 100 | Relative CFU | Low | 72 | 1.17 | 1.25 | 0.37 | 0.007 |
| 100 | Relative CFU | High | 43 | 2.33 | 2.32 | 1.34 | <0.001 |
| 100 | *Relative surf. polysacch* | *Low* | 72 | *1* | *1.18* | *1.21* | *0.917* |
| 100 | Relative surf. polysacch | High | 36 | 1.45 | 1.6 | 0.57 | <0.001 |

**Table S2.** Mutations in *mrk* operon in clones sequenced at the end of the evolution experiment.

| Environment | Strain | Ancestor | Clone ID | Gene | Mutation | Annotation |
| --- | --- | --- | --- | --- | --- | --- |
| ASM | Kva 342 | Cap + | 1A3 | *mrkD* | C→T | T26M (ACG→ATG) |
| ASM | Kva 342 | Cap + | 1A3 | *mrkD* | C→T | R291C (CGC→TGC) |
| ASM | Kva 342 | Cap + | 1A5 | *mrkB* | T→C | F8S (TTC→TCC) |
| ASM | Kva 342 | Cap + | 1A5 | *mrkD* | A→T | R41W (AGG→TGG) |
| ASM | Kva 342 | Cap - | 1D1 | *mrkF* | IS insertion | coding (345/636 nt) |
| ASM | Kva 342 | Cap - | 1D1 | *mrkC* | G→A | G439E (GGG→GAG) |
| ASM | Kva 342 | Cap - | 1D2 | *mrkD* | G→T | D36Y (GAC→TAC) |
| ASM | Kva 342 | Cap - | 1D2 | *mrkD* | C→T | S73F (TCC→TTC) |
| ASM | Kva 342 | Cap - | 1D3 | *HJIKBMFB_00805 / mrkF* | G→T | intergenic (-600/-106) |
| ASM | Kva 342 | Cap - | 1D3 | *mrkD* | C→T | R137C (CGC→TGC) |
| ASM | Kva 342 | Cap - | 1D4 | *mrkF* | G→T | Q156H (CAG→CAT) |
| ASM | Kva 342 | Cap - | 1D4 | *mrkD* | G→T | M37I (ATG→ATT) |
| ASM | Kva 342 | Cap - | 1D4 | *mrkD* | G→A | G109S (GGC→AGC) |
| ASM | Kva 342 | Cap - | 1D5 | *mrkD* | A→T | S249C (AGC→TGC) |
| ASM | Kva 342 | Cap - | 1D6 | *mrkD* | C→T | R291C (CGC→TGC) |
| ASM | Kpn BJ1 | Cap + | 1B1 | *mrkD* | T→A | S166C (AGC→TGC) |
| ASM | Kpn BJ1 | Cap + | 1B5 | *mrkD* | T→A | S151C (AGC→TGC) |
| ASM | Kpn BJ1 | Cap - | 2A1 | *mrkD* | Δ12 bp | coding (111-122/996 nt) |
| ASM | Kpn BJ1 | Cap - | 2A4 | *mrkD* | Δ12 bp | coding (111-122/996 nt) |
| ASM | Kpn BJ1 | Cap - | 2A5 | *mrkD* | T→A | S166C (AGC→TGC) |
| ASM | Kpn NTUH | Cap - | 2B1 | *mrkD* | New junction | +70bp |
| ASM | Kpn NTUH | Cap - | 2B2 | *mrkD* | C→A | G263C (GGC→TGC) |
| ASM | Kpn NTUH | Cap - | 2B3 | *mrkD* | G→A | R77C (CGC→TGC) |
| ASM | Kpn NTUH | Cap - | 2B4 | *mrkD* | G→A | S136L (TCG→TTG) |
| ASM | Kpn NTUH | Cap - | 2B4 | *mrkD* | G→A | R70C (CGC→TGC) |
| ASM | Kpn NTUH | Cap - | 2B5 | *mrkD* | C→A | G228C (GGC→TGC) |
| ASM | Kpn NTUH | Cap - | 2B5 | *mrkD* | C→A | D58Y (GAC→TAC) |
| ASM | Kpn NTUH | Cap - | 2B6 | *mrkD* | G→A | S185L (TCG→TTG) |
| LB | Kva 342 | Cap + | 2C1 | *mrkD* | Δ12 bp | coding (110-121/996 nt) |
| LB | Kva 342 | Cap + | 2C3 | *mrkD* | Δ12 bp | coding (110-121/996 nt) |
| LB | Kva 342 | Cap + | 2C5 | *mrkD* | Δ12 bp | coding (110-121/996 nt) |
| LB | Kva 342 | Cap + | 2C6 | *mrkD* | C→G | N98K (AAC→AAG) |
| LB | Kva 342 | Cap - | 3B1 | *mrkF* | Δ168 bp | coding (270-437/636 nt) |
| LB | Kva 342 | Cap - | 3B2 | *mrkD* | Δ12 bp | coding (110-121/996 nt) |
| LB | Kva 342 | Cap - | 3B4 | *mrkF* | (A)5→4 | coding (580/636 nt) |
| LB | Kva 342 | Cap - | 3B5 | *mrkD* | Δ12 bp | coding (110-121/996 nt) |
| LB | Kva 342 | Cap - | 3B6 | *mrkD* | Δ12 bp | coding (100-111/996 nt) |
| LB | Kpn BJ1 | Cap + | 2D3 | *mrkD* | Δ12 bp | coding (111-122/996 nt) |
| LB | Kpn BJ1 | Cap + | 2D4 | *mrkD* | Δ12 bp | coding (111-122/996 nt) |
| LB | Kpn BJ1 | Cap - | 3C2 | *mrkD* | C→T | G109S (GGC→AGC) |
| LB | Kpn BJ1 | Cap - | 3C2 | *mrkD* | Δ12 bp | coding (111-122/996 nt) |
| LB | Kpn BJ1 | Cap - | 3C3 | *mrkD* | Δ12 bp | coding (98-109/996 nt) |
| LB | Kpn BJ1 | Cap - | 3C4 | *mrkD* | Δ12 bp | coding (98-109/996 nt) |
| LB | Kpn NTUH | Cap + | 3A4 | *mrkD* | Δ12 bp | coding (104-115/996 nt) |
| LB | Kpn NTUH | Cap - | 3D1 | *mrkD* | (GATCCGGGGGCA)1→2 | coding (131/996 nt) |
| LB | Kpn NTUH | Cap - | 3D2 | *mrkD* | (GATCCGGGGGCA)1→2 | coding (131/996 nt) |
| LB | Kpn NTUH | Cap - | 3D3 | *mrkJ* | A→C | Q219P (CAG→CCG) |
| LB | Kpn NTUH | Cap - | 3D3 | *mrkD* | A→T | Y69N (TAT→AAT) |
| LB | Kpn NTUH | Cap - | 3D4 | *mrkD* | Δ12 bp | coding (104-115/996 nt) |
| LB | Kpn NTUH | Cap - | 3D5 | *mrkD* | New junction | coding (87/996 nt) |
| LB | Kpn NTUH | Cap - | 3D6 | *mrkD* | G→A | R57C (CGC→TGC) |
| M02 | Kva 342 | Cap + | 4A3 | *HJIKBMFB_00805 / mrkA* | G→T | intergenic (-41/-665) |
| M02 | Kva 342 | Cap + | 4A6 | *mrkH* | IS insertion | coding (95/705 nt) |
| M02 | Kpn NTUH | Cap - | 5B6 | *mrkD* | G→A | S136L (TCG→TTG) |
| AUM | Kva 342 | Cap - | 6B2 | *mrkB* | T→A | N109K (AAT→AAA) |
| AUM | Kva 342 | Cap - | 6B3 | *mrkC* | T→G | L459R (CTG→CGG) |
| AUM | Kva 342 | Cap - | 6B3 | *mrkF* | T→G | V35G (GTC→GGC) |
| AUM | Kva 342 | Cap - | 6B6 | *mrkC* | C→A | A549E (GCG→GAG) |

**Table S3.** Number of clones with mutation in mrkD.

| **Population** | **Mutation** | **Day** | **Capsulated clones with**<br>***mrkD* evolved allele** | **Non-capsulated clones**<br>**with *mrkD* evolved allele** |
| --- | --- | --- | --- | --- |
| 1A2 | Δ 119-130nt | 7 | 6/6 | 1/6 |
| 1A3 | T26M | 7 | 4/6 | 0/6 |
| 1A4 | G167C | 7 | 4/6 | 0/6 |
| 1A4 | T139P | 30 | 0/6 | 5/6 |
| 1A5 | R41W | 7 | 0/6 | 6/6 |
| 1A6 | Δ 110-121nt | 30 | 0/6 | 5/6 |

**Table S4.**
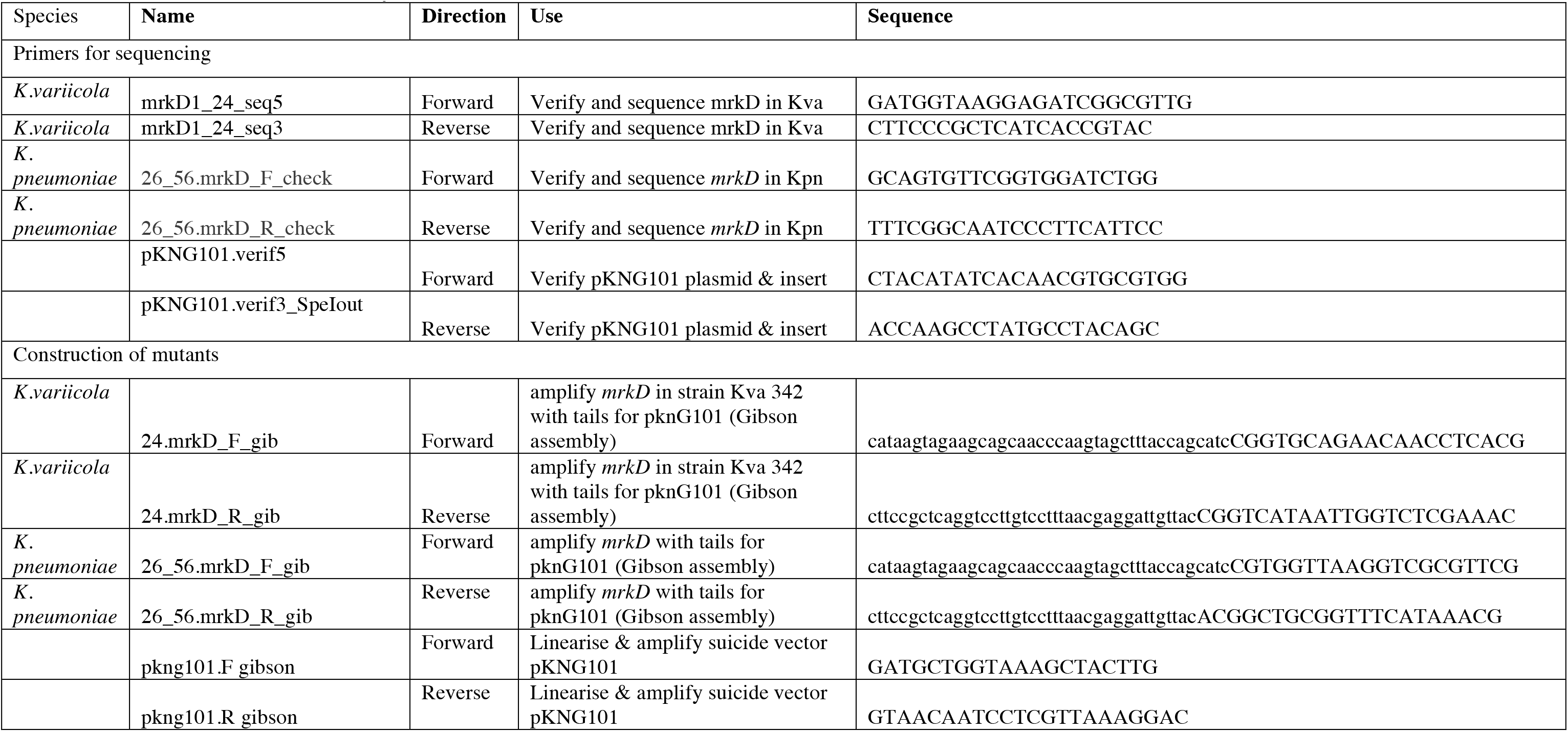
Primers used in this study.

